# Musicianship can be decoded from magnetic resonance images

**DOI:** 10.1101/2020.07.19.210906

**Authors:** Tuomas Puoliväli, Tuomo Sipola, Anja Thiede, Marina Kliuchko, Brigitte Bogert, Petri Toiviainen, Asoke K. Nandi, Lauri Parkkonen, Elvira Brattico, Tapani Ristaniemi, Tiina Parviainen

**Affiliations:** Faculty of Information Technology, University of Jyväskylä, Finland; Department of Neuroscience and Biomedical Engineering, Aalto University, Finland; Finnish Centre for Interdisciplinary Music Research, Department of Music, Art and Culture Studies, University of Jyväskylä, Finland; Department of Electronic and Computer Engineering, Brunel University London, United Kingdom; Cognitive Brain Research Unit, Department of Psychology and Logopedics, University of Helsinki, Finland; Department of Psychology, University of Jyväskylä, Finland; Centre for Interdisciplinary Brain Research, University of Jyväskylä, Finland; Center for Music in the Brain (MIB), Department of Clinical Medicine, Aarhus University & Royal Academy of Music Aarhus / Aalborg (RAMA), Denmark; College of Electronic and Information Engineering, Tongji University, Shanghai, China; Advanced Magnetic Imaging Centre, Aalto Neuroimaging, School of Science, Aalto University, Espoo, Finland

**Keywords:** Decoding, Learning, Magnetic resonance imaging, Machine learning, Musician, Support vector machines

## Abstract

Learning induces structural changes in the brain. Especially repeated, long-term behaviors, such as extensive training of playing a musical instrument, are likely to produce characteristic features to brain structure. However, it is not clear to what extent such structural features can be extracted from magnetic resonance images of the brain. Here we show that it is possible to predict whether a person is a musician or a non-musician based on the thickness of the cerebral cortex measured at 148 brain regions encompassing the whole cortex. Using a supervised machine-learning technique, we achieved a significant (*κ* = 0.321, *p <* 0.001) agreement between the actual and predicted participant groups of 30 musicians and 85 non-musicians. The areas contributing to the prediction were mostly in the frontal, parietal, and occipital lobes of the left hemisphere. Our results suggest that decoding musicianship from magnetic resonance images of brain structure is feasible. Further, the distribution of the areas that were informative in the classification, which mostly, but not entirely, overlapped with earlier findings on areas relevant for musical skills, implies that decoding-based analyses of structural properties of the brain can reveal novel aspects of musical aptitude. In particular, our results highlight differences in visual areas in addition to the already more established differences located in motor networks and networks of higher-order cognitive function.

## 1. Introduction

The gross structural properties of the brain are shaped during peri- and postnatal development, and genetic guidance continues to influence brain formation throughout infancy and childhood (see e.g. Pletikos et al., 2014). Especially during postnatal development, structural changes reflect interactions between genetic and environmental factors. However, also learning affects brain structure, and the role of the environment and individual’s own actions in influencing gray- and white-matter properties are likely to be particularly strong during development. Training-related structural changes have been observed to occur throughout the human lifespan in both the gray and white matter (Maguire et al, 2006; Scholz et al. 2009; Zatorre et al. 2012; Sampaio-Baptista et al. 2014). Clear evidence of the influence of sensory environment can be seen for example in the auditory cortices of bilingual children: more variable language and auditory environment increases the volume of the Heschl’s gyri (Ressel et al. 2010). Similarly, learning a second language increases the gray matter density of the left inferior parietal cortex in both adolescents and adults (Mechelli et al, 2004). Taken together, a magnetic resonance image (MRI) of a person’s brain should contain information about individual abilities, especially for well-trained skills that are acquired early in life.

Hence, does possessing a skill manifest as a detectable trace in the structural MRI of a person’s brain? The answer is likely to depend on at least the magnitude and type of the structural changes associated with the acquisition of a specific skill. Since playing music requires mastering of a complex set of motor and multi-sensory skills, the learning-related structural changes could be expected to be highly distributed across the cortex. Indeed, there is now a large body of evidence showing functional and structural differences between musicians’ and non-musicians’ brains (Elbert et al., 1995; Münte et al., 2002; Gaser & Schlaug, 2003a,b; Wong et al., 2007; Bermudez et al. 2009; Barrett et al. 2013; Miendlarzewska et al. 2013; Moore et al. 2014; Reybrouck & Brattico, 2015). Musicians are therefore an ideal population for testing whether the structural information obtained with MRI would allow decoding of acquired expertise, since adult professional musicians have typically started the intensive training at an early age, and it is easy to find control participants with little or no musical training.

In previous neuroscience research, machine learning has been more widely applied to functional but not so much to structural neuroimaging data. For example, studies conducted with functional magnetic resonance imaging (fMRI) have succeeded in reliably decoding reward-based decision-making processes (Hampton & O’Doherty, 2007), visual experiences and visual imagery during sleep (Nishimoto et al. 2011; Horikawa et al. 2013), moral judgments (Koster-Hale et al. 2013), and unconscious or covert mental states (Haynes & Rees, 2006). Experiments using other brain imaging modalities, namely electroencephalography (EEG) and magnetoencephalography (MEG), have demonstrated that decoding hand movement directions (Waldert et al. 2008), semantic categories of words (Cichy et al. 2014) and aware vs. unaware visual percepts (Salti et al. 2015) is possible, to name a few examples. In the specific domain of music-related expertise, it has been previously possible to decode individuals’ musicianship classes from fMRI data that was recorded while the participants were listening to different pieces of music (Saari et al., 2018). While these and other functional imaging techniques allow inferring and contrasting brain states, traits are perhaps better quantified from structural data. Indeed, functional imaging techniques only capture processes elicited by the performed task.

There are many clinical applications based on data-driven classification of structural MRIs, such as diagnosis of Alzheimer’s disease (Moradi et al. 2015), or identification of individuals at risk for bipolar disorder (Hajek et al. 2015). However, relatively few investigations on the feasibility of using machine learning techniques for classification of non-clinical MRI features have been reported to date. Given that brain structure reflects a multitude of influences and processes, and which must be relatively highly preserved to allow normal function, such tasks are inherently difficult. In this study, we show for the first time that it is possible to decode musicianship from the brain structure: to predict whether a person is a musician or a non-musician by analyzing cerebral cortical thickness data with support vector machines (SVM), which is a supervised machine-learning technique for binary classification problems (Cortes & Vapnik, 1995).

A difficult problem in structural MRI studies is identifying which findings are at the same time statistically and neurobiologically significant for a given cognitive function or behavioral skill. In principle, a small difference in one region can be much more relevant for a given cognitive function than a larger difference in another region: behaviorally (psychologically, clinically) meaningful variation is embedded within variation due to other causes across individuals. It is possible to address causal questions by disrupting brain activity for example by using transcranial magnetic stimulation, but the relative stability of brain structure renders any interventional approaches impossible in humans. Hence, how should one decide whether a statistically significant difference in cortical thickness, reported in millimeters, is meaningful in behavioral terms?

In this regard, machine-learning or decoding models may be useful: if a set of measurements systematically predicts the desired target value, one can conclude that the measurements contain information about the property of interest, which often has more value than merely reporting group differences. In this study, we attempt to decode musicianship by classifying the participants as either professional performing musicians or non-musician control participants based on their MRIs. While structural differences between musicians’ and non-musicians’ brains have been investigated for decades, it has remained unknown which cortical structures best allow decoding musicianship. By using reliable and extensively validated machine-learning and MRI processing methods, we address this issue in this study.

## 2. Materials and methods

### 2.1. Participants

A total of 121 professional, amateur, and non-musician participants were recruited to this study. Six participants were excluded from the data analysis due to neurological or psychiatric disorders. The recruited group of professionals consisted of rock, pop, jazz, and classical musicians who had started playing their first instrument before the age of thirteen. The non-musician and amateur musician groups were merged for the binary classification task. The final participant demographics are summarized to Table I. The study was part of the “Tunteet” protocol approved by the Coordinating ethics committee of the Helsinki and Uusimaa Hospital District. Written informed consent was obtained from each participant prior to the measurements. The research was performed in compliance with the Declaration of Helsinki by World Medical Association.

**Table 1:** Participant demographics. Handedness missing for two participants.

| Group | N | Age |  |  | Gender |  | Handedness |  |  | Years training |  |
| --- | --- | --- | --- | --- | --- | --- | --- | --- | --- | --- | --- |
|  |  | Mean | SD | Range | Male | Female | Left | Right | Ambidext. | Mean | SD |
| Non-mus | 80 | 28.61 | 8.15 | 19–53 | 35 | 50 | 6 | 78 | 1 | 2.15 | 3.81 |
| Musician | 35 | 27.53 | 7.19 | 18–45 | 16 | 14 | 2 | 28 |  | 13.00 | 7.62 |
| Total | 115 |  |  |  | 51 | 64 | 8 | 106 | 1 |  |  |

### 2.2. MRI data acquisition

The MR images were acquired at the Advanced Magnetic Resonance Imaging (AMI) center of Aalto Neuroimaging, Aalto University (Espoo, Finland). A Siemens Magnetom Skyra 3 T whole-body scanner (Siemens Healthcare, Erlangen, Germany) with a standard 20-channel head-neck coil was used. A gradient-echo (MP-RAGE) T1-weighted sequence with repetition time, echo time, inversion time, and flip angle of 2530 ms, 3.3 ms, 1100 ms, and 7 degrees, respectively, was used. Voxel size was set as 1 mm^3^.

### 2.3. Extraction of cerebral cortical thickness data

Automated cerebral cortical thickness measurements were performed using the FreeSurfer MRI analysis software suite (Dale et al. 1999; Fischl et al. 1999), which is available online at https://surfer.nmr.mgh.harvard.edu/. FreeSurfer has been shown to make accurate and precise volumetric estimates in several validation studies (Fischl et al. 2002; Han et al. 2006; Wonderlick et al. 2009; Cardinale et al. 2014). The left and right hemispheres of the cerebral cortex were both parcellated into 74 regions according to the Destrieux atlas (Destrieux et al. 2010). The average thickness of gray matter at each of the 148 regions was estimated. The intermediate steps of the automated procedures were manually checked for obvious errors, and corrected, if needed. The smallest measured cortical thickness was 1.47 mm and the largest 4.14 mm, suggesting that the data did not contain any major outliers, since the human cerebral cortical thickness is known to vary approximately between 1.0 and 4.5 mm (Fischl et al. 2000).

### 2.4. Control for confounding effects

Cortical thickness decreases with age (Salat et al. 2004; Thambisetty et al. 2010; Toga et al. 2011). Since the participants in our study spun a wide range of ages, the effect may have consequences for our study. If, for example, we trained a classifier with young musicians, and tested it with old musicians, the performance would be probably worse than with age-balanced groups. To test the existence of this effect in our data, we fitted a regression model using age as the independent variable and mean cortical thickness as the dependent variable, and examined the coefficient of determination. A successful replication of the previous studies would imply a need for age-balanced training and test sets. Because previous studies have also identified structural differences between the brains of males and females (Im et al. 2006; Sowell et al. 2007), we also examined the mean cortical thickness with respect to the participants’ gender.

### 2.5. Classification and cross-validation

Supervised machine learning is about finding a mapping *X → Y* where *x*_*i*_ ∈ *X* is a sample and *y*_*i*_ ∈ *Y* is a group label, when having a limited number of labeled samples (*y*_1_, *x*_1_), …, (*y*_*n*_, *x*_*n*_). Using SVMs (Cortes and Vapnik, 1995), we studied a two-class problem with *y*_*i*_ ∈ { musician, non-musician and each sample *x*_*i*_ containing the cerebral cortical thickness measurements of the participant *i*. The SVM classifier was selected because it is known to perform well in many kinds of classification tasks (Fernández-Delgado et al. 2014) and its implementation is available in many machine-learning packages.

To use the data as efficiently as possible, we used nested stratified k-fold cross-validation for model selection and validation (Kohavi, 1995) (Figure 1). We selected to test linear and radial basis function kernels for the SVM and k = 10 for both the inner and outer cross-validation loops. Since age but not gender was identified to have an effect on the mean cortical thickness (see Sections 3.1 and 3.2), we balanced the folds with respect to participant and age groups. The coarse logarithmic grid { 1, 10, 100, 1000} was searched to optimize the SVM *C* parameter, which controls the generalization ability. Since the participant groups were imbalanced, the parameter *C* was adjusted for group *j* as *Cw*_*j*_ where *w*_*j*_ is a weight inversely proportional to the group frequency.

**Figure 1:**
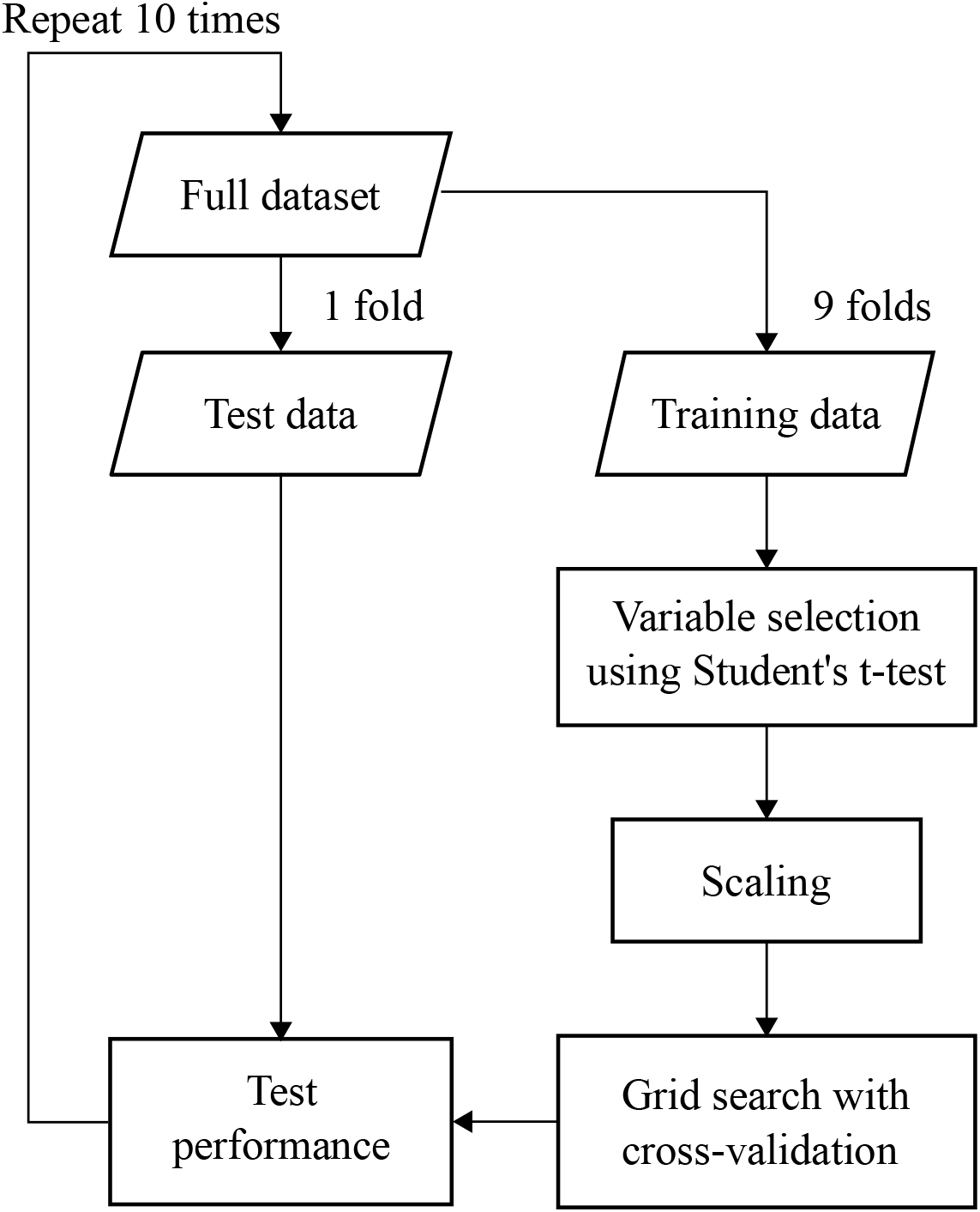
Nested cross-validation using ten-fold cross-validation as the outer loop. The model was constructed using 9 folds of the data and tested with the remaining one. The process was repeated 10 times, so that each fold acted as the test data once. The SVM parameters were selected using a grid search with a second cross-validation loop.

The machine learning pipeline was implemented in Python version 3.2.0 from the Python Software Foundation. For cross-validation and classification we employed the Scikit-learn module (Pedregosa et al. 2011; Abraham et al. 2014), which internally uses the LIBSVM library (Chang and Lin, 2011) for SVMs. The cortical surface visualizations were produced with PySurfer, which is available online at https://pysurfer.github.io/.

### 2.6. Variable selection and scaling

To identify an informative subset of the most relevant variables for constructing the classifier, we used a filter-type variable selection procedure as a preprocessing step (Weston et al., 2000; Gyuon & Elisseef, 2003). The Student’s t-test for two independent samples with the critical level *α* = 0.05 was used to discard variables for which the null hypothesis of equal means could not be rejected. The variable selection was repeated on each cross-validation iteration. Therefore, we used the number of times a variable was selected as an indicator of its relative importance. The selected variables were transformed to standard scores by removing the mean and scaling to unit variance to avoid variables with large numerical ranges to dominate the classifier fitting process.

### 2.7. Evaluation of classifier performance

We used Cohen’s kappa (Landis and Koch, 1977) as the primary metric for evaluating the classifier’s performance. This metric is zero for chance-level agreement between the actual and predicted participant groups, +1 for complete agreement, and *−*1 for complete disagreement. It is calculated as *κ* = (*π*_*o*_ *−π*_*e*_)*/*(1*−π*_*e*_) where *π*_*o*_ is the observed agreement (percentage of correct predictions) and *π*_*e*_ is the probability of an agreement occurring by chance. The metric can be rewritten as *κ* = 1 *−* (1 *− π*_*o*_)*/*(1 *− π*_*e*_), where 1 *− π*_*o*_ is the error rate and 1 *− π*_*e*_ is the probability of a disagreement occurring by chance.

### 2.8. Statistical assessment of over-fitting

To investigate the reliability of the classifier, we used the following permutation test. First, the group labels of the entire data were randomly permuted. The classifier training and testing were then performed with the expectation of achieving chance-level prediction metrics on average. Next, this process was repeated 1000 times and the prediction metrics collected as the null distribution. Finally, the classifier’s performance was compared to this distribution. If the probability of obtaining equal or better results with the randomly-permuted group labels was less than the critical level *α* = 0.05, we considered the classifier reliable.

## 3. Results

### 3.1. Mean cortical thickness as a function of age

The relationship between the participants’ age and mean cerebral cortical thickness appeared to be linear; see Figure 3 for an illustration. A regression analysis showed that the mean cortical thickness decreased significantly with age (*F*_1,114_ = 39.72, *p <* 0.001); mean cortical thickness = 2.652 *−*0.005 *×*age; *r*^2^ =0.260. The 95 % confidence interval for the slope was [–0.007, –0.003]. The result was in agreement with previous studies (Salat et al. 2004; Thambisetty et al. 2010; Toga et al. 2011) and implied a need for age-balanced cross-validation folds.

### 3.2. Mean cortical thickness as a function of gender

The mean cerebral cortical thickness was 2.51 mm (standard deviation was 0.90 mm) for the male participants and 2.51 mm (0.70 mm) for the female participants. The 95 % confidence intervals of the means were [2.26, 2.75] and [2.34, 2.68] for males and females respectively. The measurements were approximately normally distributed in both groups. There was no significant difference between the means according to an independent two-sample t-test: *t*(113) = 0.036, *p* = 0.971. The data are illustrated with box plots in Figure 2. The result implies that the cross-validation folds do not need to be balanced with respect to gender.

**Figure 2:**
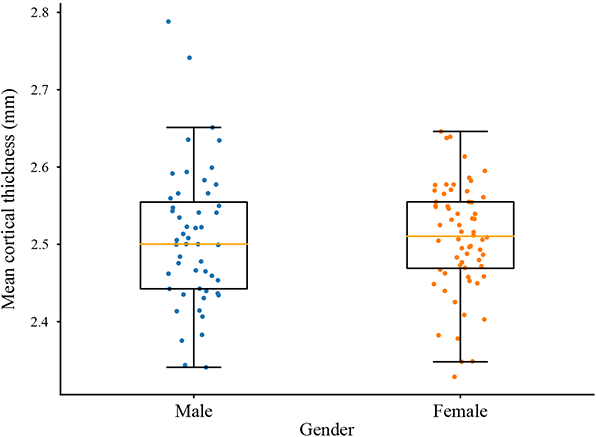
Mean cerebral cortical thickness (mm) as a function of age (years). Age explained 26 % of variation in the mean cortical thickness.

**Figure 3:**
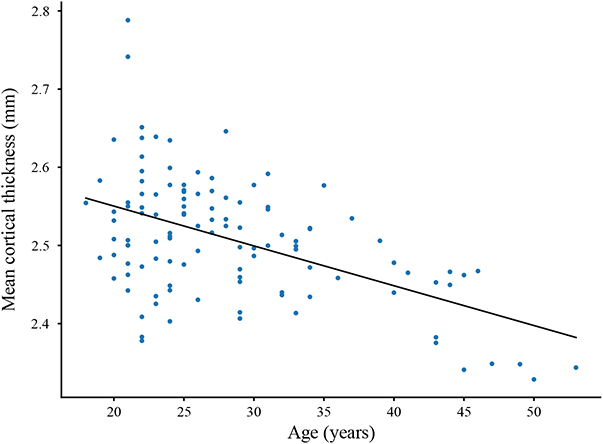
The mean cerebral cortical thickness was equal in males and females.

### 3.3. Classifier performance

The linear SVM with nested cross-validation and group-frequency-weighted samples achieved a significant inter-rater agreement (*κ* = 0.321, *p <* 0.001) between the actual and predicted participant groups which was an imbalanced sample of musicians and control participants. It correctly classified 19 of the 30 (63.3 %) musician and 62 of the 85 (72.9 %) non-musician participants. In the set of 73 participants classified as non-musicians, 62 (84.9 %) were actually non-musicians. In the set of 42 participants classified as musicians, 19 (45.2 %) were actually musicians. The confusion matrix of the classification results is shown in Table II. A radial basis kernel function for the SVM was also tested, but the results were not analyzed further due to over-fitting, which was assessed using the permutation test described in section 5.8 (*p >* 0.05).

**Table 2:** SVM classification results. Sample size was *N* = 115 participants.

|  | Non-musician | Musician | Total |
| --- | --- | --- | --- |
| Non-musician | 53.9 % | 20 % | 73.9 % |
| Musician | 9.6 % | 16.5 % | 26.1 % |
| Total | 63.5 % | 36.5 % | 100 % |

Post-hoc tests were performed to assess possible causes of participant misclassification. For the demographic data, there was a significant difference in age between the correctly and incorrectly classified non-musicians (Student’s t-test; *p <* 0.005) such that incorrectly classified non-musicians were older than correctly classified non-musicians. For the imaging data, there was a significant difference in mean cortical thickness between the incorrectly and correctly classified non-musicians (Student’s t-test; *p <* 0.001). The same effect was observed for musicians (Student’s t-test; *p <* 0.005). Given that cortical thickness was smaller in musicians than in non-musicians in the frequently selected regions of the machine learning pipeline (see section 3.5), and that cortical thickness decreased with age (see section 3.1), this suggests that sometimes the classifier confused the effects of musical training and aging because cortical thinning is associated with both of these two processes in these data.

### 3.4. Assessment of over-fitting with permutation tests

The classifier training and testing was repeated 1000 times with randomly permuted group labels. The mean inter-rater agreement was *κ*_*µ*_ = *−*0.002 and the standard deviation was 0.124. The 95 % confidence interval for the mean inter-rater agreement was [*−*0.010, 0.006]. The permutation distribution was approximately normally distributed. The probability of observing a Cohen’s kappa equal to or larger than *κ* = 0.321 by chance was *p* = 0.006. The permutation test results are illustrated as a histogram in Figure 4. The results suggest that the actual classifier was reliably fit to the data and not prone to over-fitting, since it did not perform well with the randomly permuted group labels.

**Figure 4:**
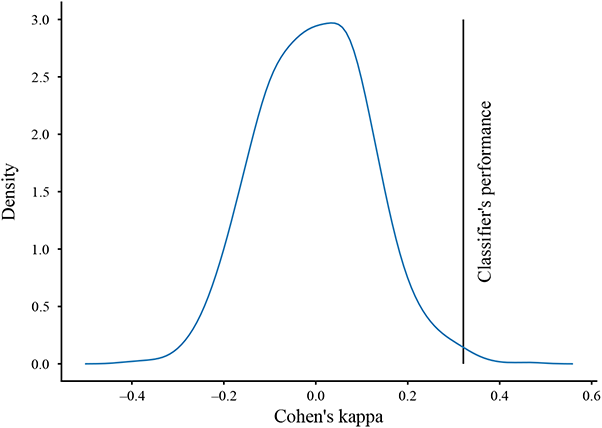
The permutation-test results over 1000 iterations. The density estimate shows the distribution of the Cohen’s kappa statistic when the group labels “musician” and “non-musician” were randomly permuted and the machine learning repeated. The black vertical bar indicates the actual classifier performance.

### 3.5. Variable selection

The variable selection results of the linear SVM are summarized to Table III and illustrated in Figure 5. The results were fairly stable, meaning that similar decisions were made on each of the ten cross-validation iterations. There was a total of thirteen regions, nine in the left hemisphere and four in the right hemisphere, which were selected on every iteration. Five more regions, all located in the left hemisphere, were selected on more than half of the iterations. In total, 39 different regions were selected during the ten cross-validation iterations. Because it is difficult to assess the importance or meaning of those regions that were selected infrequently, we chose to analyze further only those 18 regions that were selected on more than half of the iterations.

**Table 3:** Variable selection results

| Region | Lobe | # | Musicians | Non-mus | Diff |
| --- | --- | --- | --- | --- | --- |
| **Left hemisphere** |  |  |  |  |  |
| Middle frontal gyrus | Frontal | 10 | 2.57 | 2.63 | -0.060 |
| Middle frontal sulcus | Frontal | 8 | 2.165 | 2.218 | -0.053 |
| Paracentral lobule and sulcus | Parietal | 10 | 2.285 | 2.38 | -0.095 |
| Intraparietal sulcus and trans. parietal sulci | Parietal | 10 | 2.118 | 2.2 | -0.072 |
| Angular gyrus | Parietal | 9 | 2.608 | 2.679 | -0.071 |
| Superior parietal lobule | Parietal | 7 | 2.369 | 2.424 | -0.055 |
| Posterior transverse collateral sulcus | Occipito-temporal | 10 | 2.003 | 2.115 | -0.112 |
| Cuneus | Occipital | 10 | 1.817 | 1.891 | -0.074 |
| Middle occipital gyrus | Occipital | 10 | 2.498 | 2.592 | -0.094 |
| Superior occipital gyrus | Occipital | 10 | 2.144 | 2.245 | -0.101 |
| Superior occ. sulcus and transv. occ. sulcus | Occipital | 10 | 2.033 | 2.12 | -0.087 |
| Anterior occipital sulcus and preoccipital notch | Occipital | 10 | 2.188 | 2.273 | -0.085 |
| Middle occipital sulcus and lunatus sulcus | Occipital | 8 | 1.974 | 2.035 | -0.061 |
| Parieto-occipital sulcus | Occipital | 9 | 2.189 | 2.262 | -0.073 |
| **Right hemisphere** |  |  |  |  |  |
| Angular gyrus | Parietal | 10 | 2.675 | 2.752 | -0.077 |
| Parieto-occipital sulcus | Parieto-occipital | 10 | 2.21 | 2.299 | -0.089 |
| Medial occ.-temp. sulcus and lingual sulcus | Occipito-temporal | 10 | 2.417 | 2.485 | -0.065 |
| Inferior occipital gyrus and sulcus | Occipital | 10 | 2.506 | 2.613 | -0.107 |

**Figure 5:**
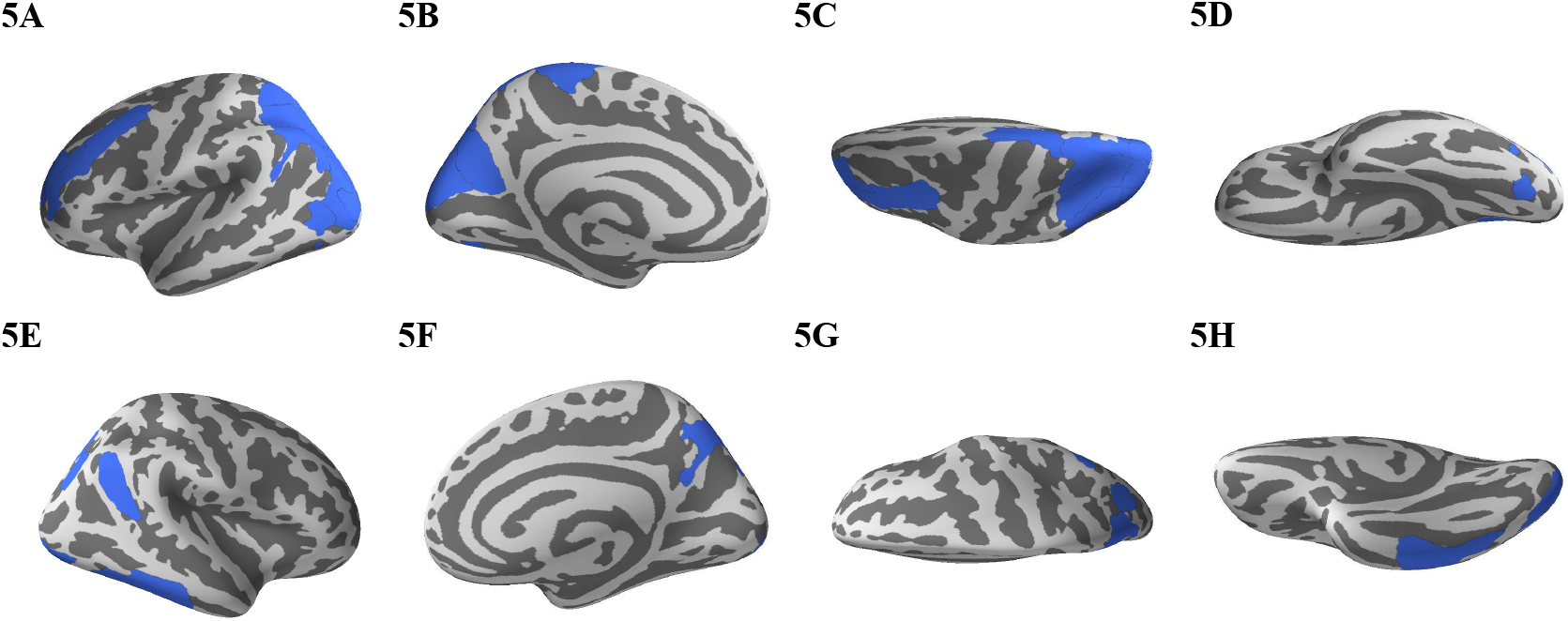
The feature selection results of the linear SVM. The regions that were selected on more than half of the cross-validation iterations are shown in blue. **A** lateral view of the left hemisphere; **B** medial view of the left hemisphere; **C** dorsal view of the left hemisphere; **D** ventral view of the left hemisphere; labels from **E** to **H** show the right hemisphere.

The frequently selected regions were located in the frontal, parietal, and occipital lobes. Since these feature selection results themselves did not contain information about the direction or magnitude of effects, we compared the cortical thickness in the frequently selected regions between musicians and non-musicians as a post-hoc comparison in order to understand the effects of any underlying musical ability better. The results are presented in Table III. The cortical thickness of every frequently selected region was smaller in musicians than in non-musicians. To have a reference for the effect sizes in these regions, one can compare the rate of cortical thinning with aging to the differences observed between musicians and non-musicians. The rate of cortical thinning was 0.005 mm per year on average in our data (*p <* 0.001, see results in Section 3.1). Therefore, the differences between the musicians’ and non-musicians’ brains were approximately 10–20 times larger than yearly cortical thinning.

## 4. Discussion

Our results demonstrate that decoding musicianship from brain MRIs is feasible. The linear SVM achieved a significant inter-rater agreement between the actual and predicted participant groups. In other words, it was able to learn a function from labeled MRI data that could be used to classify new previously unseen unlabeled MRI data. It correctly labeled 63.3 % of the musician and 72.9 % of the non-musician participants.

Since playing music is a combined multi-sensory and motor skill, we expected that the relevant regions for classifying the participants would be widely distributed across the cortex. Indeed, the classifier’s ability to distinguish musicians from non-musicians was based on differences in cortical thickness in several regions located in the frontal, parietal, and occipital lobes with a greater emphasis on the left compared to the right hemisphere. This is in general agreement with the previous literature, which demonstrates that several areas in frontal, parietal and occipital lobes are linked to musical ability.

In line with our results, James and colleagues (2012) found a decreased gray matter density in amateur and professional musicians compared to non-musician participants in the right postcentral gyrus, bilateral paracentral lobule, bilateral precuneus, left inferior occipital gyrus, and bilateral striatal areas. Our results are in line with these findings especially for the medial surface regions, however suggesting more emphasized role for the left than right hemisphere. In our study, the thickness of the left precuneus and the left paracentral lobule were observed to be relevant predictors of musical ability. James and colleagues (2012) interpreted their findings to reflect automatized sensorimotor function and argued that fewer neurons are eventually needed for producing the playing-related movements, after a certain level of proficiency has been acquired, which could explain also our results.

Indeed, increases in proficiency during development are often associated with decreases in gray matter, and as musical skills are often acquired early in life, it would make sense that the areas that are engaged in increased amounts of sensorimotor training show stronger expertise-related pruning. The volume of left precuneus, which is a part of the superior parietal lobule, and left paracentral lobule were observed as relevant predictors of musical ability both by James and co-authors and the current study, highlighting the role of these regions in acquired musical aptitude. James and colleagues (2012) found also regions with increased gray-matter densities but they did not overlap with the regions we observed to be important. Achieving detailed understanding on the differences between expertise-related decrease vs. increase in gray matter thickness and volume would require analysis at the level of underlying neuronal and vascular changes. It may well be, that the skill acquisition during development builds on very different neuroplastic processes which are integrated with maturational changes, than the skill acquisition later in life (Steele et al., 2012; Vaquero et al., 2015; de Manzano and Ullén, 2017; Weisberg et al., 2019).

Further, while many studies that investigated skill- and experience-related changes in the human brain structure report increases in volume, area, or thickness, the large amounts of knowledge, experience, and skills obtained during a lifetime are ultimately not likely to be represented in the brain by cumulative, continual increases in structural measures (Wenger et al., 2017; Calmels et al., 2019). Indeed, selection, deletion and pruning are effective mechanisms that occur throughout the nature, including brain development. The expansion-renormalization model posits that learning is a two-step process, in which the first step involves an expansion of gray matter structure, followed by selection and pruning in the second step, thus keeping the overall brain volume approximately constant (Wenger et al., 2017). Since musical expertise is acquired over very long time scales, these data could be explained in the context of the expansion–renormalization model by a gradually accumulated negative net effect, which is either due to over-pruning over time, declined expansion volumes over time, or both. These processes might be modulated by the already acquired experience or aging.

Gray-matter differences in the frontal lobe between musicians and non-musicians have also been reported in previous experiments. Gaser & Schlaug (2003a) found an increased gray-matter volume in the left inferior frontal gyrus in musicians. Another study (Bermudez et al. 2009) observed greater cortical thickness bilaterally in the middle frontal gyrus, and the left superior frontal gyrus. Our results are partially in conflict with these observations, as we detected greater cortical thickness in the left middle frontal gyrus and sulcus in non-musicians than musicians. Unfortunately, it is difficult to point out what could be the reason for the disagreement due to differences in participant recruitment and data analysis methods. While Bermudez and colleagues performed voxel-level analyses based on the general linear model, we used supervised machine learning and analyzed the data using a predefined cortical parcellation.

Our findings on the significance of occipital areas for classifying musicians vs. non-musicians may seem at first contradictory, as playing an instrument is strongly linked with auditory-motor-integration. There are however also earlier studies showing this association. Luders and colleagues (2004) identified structural changes in the occipital lobe related to musical ability. They found gray matter asymmetries in musicians in the cuneus and the medial occipital gyrus. Along the same lines, Bengtsson and co-workers (2005) showed that practicing playing music during adolescence correlated with fractional anisotropy in the splenium and body of the corpus callosum extending into the white matter of the occipital lobe. Foster and Zatorre (2010) found a positive correlation between a relative pitch task and cortical thickness in the left precuneus, left inferior parietal lobule, bilateral parieto-occipital sulcus, left middle frontal gyrus, left precentral sulcus, right precentral gyrus, and left ventrolateral frontal cortex. A recent fMRI study also indicated large increases in brain-scale networks connected to music-selective visual areas (Mongelli et al., 2017).

The observed cortical differences between musicians and non-musicians in the occipital lobe could be related to an audiovisual or visuo-motor integration process, that presumably forms an essential component in learning to fluently translate visual notes to auditory and motor representations: Interestingly, modulating attentional demands for auditory stimuli has been shown to activate the peripheral visual cortex not only in blind subjects but also in normally sighted adults (Cate et al. 2009). The cross-modal sensory interactions might thus be more readily available for relevant behavioral needs than is typically assumed (Shimojo & Shams, 2001). Music training might well be one of the skills that readily maintains integration between auditory, visual, and motor cortices, at the early level of cortical hierarchy. Indeed, previous studies have found musicians to score better than non-musicians in visual attention tests (Rodrigues et al. 2013), as well as in audiovisual timing tests where musicians were better in judging whether auditory and visual stimuli were presented synchronously (Lu et al. 2014). Moreover, an EEG study found that musicians are able to expect auditory endings based on visually-presented musical sequences (Schön and Besson, 2003).

As with any comparison between groups differing in the level of expertise, it is difficult, if not impossible, to tease apart the influence of expertise and innate differences in brain structural properties. Most of the previous studies investigating the neural basis of musical expertise have relied on cross-sectional experimental designs. A potential problem shared by such studies, including the present one, is that they are able to rule out only certain confounding effects. Therefore, it is interesting to compare the results on relevant cortical areas for musical expertise between cross-sectional and longitudinal or intervention studies. For example, Hyde and colleagues (2009) studied the effect of fifteen months of musical training in musically untrained children, compared to control children receiving no training. The children who received the musical training showed a decrease in the relative volume of the left middle occipital gyrus, which was a relevant predictor of musical ability in our study, too. These common findings give rather strong evidence of the importance of this region in acquiring musical aptitude. Interestingly, learning to play a musical instrument is a task that first relies much on visual feedback for achieving the desired motor function (Groussard et al. 2014). Early sensory areas show also the fastest developmental trajectory. Therefore, it is not surprising that the region that was found relevant for musical aptitude in both studies was located in the occipital lobe.

Hyde and colleagues (2009) also found increased relative volumes in frontal and temporal lobes but we did not observe such effects. The disagreement could be related to the participants’ age, since Hyde and colleagues studied young children whereas we investigated adults. Indeed, the cortical gray matter shows considerable changes, both increase and decrease, in thickness and volume during protracted period of development (Toga et al, 2006). These developmental processes continue well into adolescence and even adulthood for some brain regions, and reflect the combined influence of genetic and environmental factors. The discrepancy might also stem from the very different duration of received musical training; our musician participants had practiced their instrument thirteen years on average.

To make the classifier generalize optimally, we investigated the effects of age and gender on the participants’ mean cortical thickness. Since an analysis of the entire sample of musicians and non-musicians before training the classifier would lead to over-fitting, the effects of age and gender were studied without knowledge of participant groups. The mean cortical thickness decreased with age, which replicated previously reported results (Salat et al. 2004; Thambisetty et al. 2010; Toga et al. 2011). While some studies have also reported cortical differences between males’ and females’ brains (Im et al. 2006; Sowell et al. 2007), we did not observe such an effect on the level of the whole cortex. Therefore, the participants’ group and age – but not gender – were used in balancing the classifier’s cross-validation folds. It should be noted that since age and the number of years of experience in playing music are naturally correlated in professional musicians, a limitation in this study is that it is not possible to disentangle all interactions between development, learning, and aging.

Our feature-selection results indicate that the left-hemispheric regions were more informative than the right-hemispheric regions for classifying the participants as either musicians or non-musicians; only four reliable regions got selected from the right hemisphere, and as many as 14 from the left hemisphere (see Table III). Thus, future studies could benefit from expanding the feature set by calculating suitable inter-hemispheric lateralization scores, since differences in hemispheric asymmetry between musicians and non-musicians have been reported in previous studies (Schneider et al. 2005; Ellis et al. 2013; Burunat et al. 2015). However, this imbalance could be alternatively partially related to sampling; most of our participants were right-handed.

Remarkably, the temporal regions of the Destrieux atlas were not considered informative by our feature-selection procedure, which was based on cross-validation and the Student’s t-test. However, this may be simply due to the relatively large sizes of the Destrieux atlases temporal regions resulting in the loss of information regarding musical ability.Given the availability of many different cortical parcellations (Fischl et al., 2004; Fan et al., 2016; Glasser et al., 2016; Gordon et al., 2016; Schaefer et al., 2018) which are based on different delineation methods and region sizes, alternative choices can be tested in future studies. It is also possible that cortical area or volume would reflect training-related changes better than cortical thickness, which can be also tested in subsequent studies. Indeed, cortical thickness and area are known to be influenced by different sets of genetic factors and neuroplastic processes (Panizzon et al., 2009; Raznahan et al., 2011; Walhovd et al., 2016). In addition, thickness and area also have different developmental trajectories and are generally differentially associated with cognitive abilities (Vuoksimaa et al., 2016; Winkler et al., 2017). Taking the opposite view, the present results allow speculation on regions that are critical for identifying professional musicians. It may be that fine-tuned auditory sensory function is not sufficient for separating musicians from non-musicians. In fact, sensory expertise might be generally much more balanced over different sub-populations than motor expertise. Professional musicians must master fine motor movements and repertoires and be efficient in integrating visuo-motor functions (and memory) with auditory perception, which might cause more unique structural patterns that become represented in musicians’ brains. Thus, the brain networks related to actions performed by musicians might be much more discriminative than the brain networks related to perception.

On a technical and more general level, it is of importance also to consider the advantages and disadvantages of using a brain-atlas-based analysis approach in the first place. On one hand, a voxel-level analysis would allow detecting very focal effects. However, as the number of voxels would be orders of magnitude larger than the number of brain regions in a typical atlas, much more emphasis would be required on handling over-fitting and multiple testing in particular, which in turn would usually imply a need for obtaining a larger sample of participants. On the other hand, brain atlases allow making neuroanatomical definitions, communicating findings easily, and facilitate comparisons across studies. Ideally, the regions of a brain atlas also delineate structures with specific functions, allowing efficient data interpretation. Disadvantages of using brain atlases have been recently reviewed by Dickie and colleagues (2017) and include, for example, experimental effects being localized between regions and risk of systematic biases. Finally, it should be noted that voxel-based analyses typically require the data to be smoothed substantially, which can make the actual effective resolution close to using a very fine-grained brain atlas. From this perspective, these two approaches can be fairly similar with particular choices of parameters.

In this study, we chose to decode musicianship from MRIs. Since this was shown to be feasible, as a next step it would be interesting to decode more specific musical skills. For example, it might be possible to decode musicians’ main instruments using a multi-class classification method. In such approach, several one-vs-one or one-vs-many SVMs could be used (Hsu & Lin, 2002). Here, while our sample did include musicians with different main instruments, the sample size of thirty professional musicians would have been too small to allow analysis of sub-groups. Another way to make the classification problem more fine-grained could be to predict a continuous value such as years of practice instead of musicianship. Including both structural and functional features in such decoding analyses (see e.g. Albouy et al., 2019) would be particularly interesting, since it would become possible to assess whether particular structural differences are associated with corresponding functional differences and how such relationships change as a function of (or type of) accumulated experience.

Overall, our data are consistent with the previous studies that suggest professional musicians’ brains to be structurally different compared to non-musicians’ brains. Here we demonstrated that using SVMs, such differences can be detected from MRIs on the level of individual participants. While the most plausible explanation is that the observed differences are caused by the decades-lasting intensive musical training, it is impossible to make a definite statement since the musicians’ brains could be different to begin with; Merrett and colleagues (2013) have recently reviewed possible confounding variables such as genetics, personality, and early auditory environment. Similarly, we are not able to rule out transfer effects from other multi-sensory and motor skills associated with musical ability. In future studies, it should be considered if behavioral data could be included as covariates (e.g. data from an auditory or motor task in which musicians could be assumed to perform better). Studies investigating training- or learning-induced structural changes in the brain have also been criticized for small effect sizes compared to imaging accuracy (Thomas & Baker, 2013). However, it should be acknowledged that not all structural changes are necessarily created equal: for a given behavioral function small structural differences could sometimes be more relevant than larger ones. In our study, the relatively large sizes of the analyzed regions emphasize differences in cortical thickness that occur consistently over the defined regions; very focal changes could become insignificant in the average that is computed from each region. Further, the novelty in our approach is that in addition to finding which brain regions are different between musicians and non-musicians, we also obtain an estimate of how important those regions are for decoding musicianship.

Future studies should explore ways of improving the classification accuracy. The performance of SVMs could be compared to other available classification methods such as random forests or neural networks. Different feature selection methods could be also tested. In particular, it would be interesting to contrast filter and wrapper type methods (Weston et al., 2000). Increasing the sample size or MRI resolution (e.g. performing imaging at 7 T) could also improve the results.

## 5. Conclusions

In this study, we have demonstrated that it is feasible to decode musicianship from structural MRIs of the brain. Using well-known and previously extensively validated machine learning and MRI processing techniques, we analyzed a fairly large sample of 30 professional musician and 85 non-musician control participants. We were able to correctly categorize 63.3 % of the musician and 72.9 % of the non-musician participants based on cerebral cortical thickness. The ability to distinguish the musicians from the non-musicians was based on the thickness of several regions located mostly in the frontal, parietal, and occipital lobes of the left hemisphere. While several previous studies have compared structural MRIs of musicians’ and non-musicians’ brains, it has remained unclear how relevant the revealed differences are for musical aptitude. In addition to simply identifying differences in the brains of musicians and non-musicians, we could estimate how important they are in predicting a person’s musical ability. Overall, our results suggest that predicting musicianship from brain structure is a challenging task but solvable to a certain degree.

## Acknowledgements

This research was financially supported by TEKES of Finland under the grant 40334/10 “Machine learning for future music and learning technologies”. We thank IT Center for Science Ltd, administered by the Finnish Ministry of Education and Culture, for providing data storage and computing resources. We also thank Ilkka Pölönen and Jarmo Hämäläinen for helpful discussions, Jussi Numminen for performing a medical screening of the collected MRIs, and Jyrki Mäkelä for contributing to initial stages of the project and for supervision of participant recruitment. We thank Marita Kattelus and Toni Auranen for helping in the data acquisition. A. K. Nandi thanks TEKES for the award of Finland Distinguished Professorship.

